# Alignment-free Bacterial Taxonomy Classification with Genomic Language Models

**DOI:** 10.1101/2025.06.27.662019

**Authors:** Mike Leske, Jamie A. FitzGerald, Keith Coughlan, Francesca Bottacini, Haithem Afli, Bruno Gabriel Nascimento Andrade

## Abstract

Advances in natural language processing, including the ability to process long sequences, have paved the way for the development of Genomic Language Models (gLM). This study evaluates the feasibility of four models for bacterial classification using 16S rRNA sequences and demonstrates that gLM embeddings can be applied to effectively classify sequences at the species level, matching or outperforming the accuracy of established bioinformatics tools like BLAST+ and VSEARCH. We adopt *cosine similarity* as a computationally efficient metric, enabling classification orders of magnitude faster than current methods, and show that it carries biologically relevant signals. In addition, we demonstrate how sequence embeddings can be used to identify mislabeled sequences. Our findings place gLM embeddings as a promising alternative to traditional alignment-based methods, especially in large-scale applications such as metataxonomic assignments. Despite its wide potential, key challenges remain, including the sensitivity of embeddings to sequences of different lengths.

## 1 Introduction

The advent of high-throughput sequencing technologies has revolutionized our understanding of microbial communities by enabling comprehensive analyses of their composition and function. A cornerstone of these studies is the 16S ribosomal RNA (rRNA) gene, a highly conserved component of the prokaryotic ribosome containing variable regions serving as unique ‘molecular fingerprints’, allowing microbes to be distinguished and classified based on sequence similarities [1–4]. Sequencing and analysis of the 16S rRNA gene have become standard practice in identifying and classifying bacteria, often down to the genus or species level. This approach plays a critical role in metagenomic studies by allowing the estimation of the abundance and diversity of bacteria within complex ecosystems. The field of genomics has recently witnessed the emergence of Genomic Language Models (gLMs), including Evo [5], Nucleotide Transformer [6], and DNABERT-2 [7], which apply principles from Natural Language Processing (NLP) to DNA sequences. These models consider genomic sequences as a language, capturing the contextual relationships between nucleotides to predict functional attributes of the genome. Recent studies have demonstrated the potential of gLMs in various applications, including motif and fitness prediction, host classification, and sequence design and generation, underscoring their ability to improve our understanding of genomic data [8].

Since the development of the Transformer architecture [9], the NLP domain has seen unprecedented progress. By processing whole text sequences in parallel through Transformer blocks, a computational (self-)attention mechanism allows the model to learn complex word interactions and attain a complete “language model”. Transformer-based (large) language models like BERT [10], GPT-2/3/4/o1 [11–13], Gemini [14], the Llama model family [15, 16], or DeepSeek [17] have benefited from parallel GPU processing and the global accumulation of digital text data across the internet. Today, Large Language Models are trained on trillions of words and are made of (hundreds of) billions of parameters. Typical language model architectures include *encoder-only, decoder-only*, and *encoder-decoder. Encoder-only* models transform input text into numerical representations and are often used for text understanding tasks, such as classification or sentiment analysis. These numerical representations are commonly known as *embeddings*. The process of obtaining expressive numerical representations of input sequences is commonly known as Representation Learning [18]. *Decoder-only* models learn an internal representation of the input text that supports the model in predicting the most likely next token (character, syllable, or word), thus generating an output that is indistinguishable from human-generated text. *Encoder-decoder* models excel in translation tasks, where an input sequence is translated into an output sequence.

Similarly to natural languages, the genome is also a language itself, composed of four nucleotide letters, which combine into genes (words) whose interactions compose functional pathways (phrases). Similarly to a language, syntactic (molecular) rules define how genes should interact with each other. As the architecture of Transformer models improved to process sequences of gradually increasing length, a new field of Genomic Language Models has emerged in which Transformers and their potential successor technologies are trained on massive genome datasets. The ability to process extremely long sequences compared to natural Language Models is required as bacterial and eukaryotic genomes are multiple million base pairs long, and gene regulation often depends on very distant regions of the genome. Some of the most prominent Genomic Language Models applied to date are DNABERT-2, DNABERT-S [19], Nucleotide Transformer (all *encoder-only*), and Evo (*decoder-only*), which have been used for sequence classification, promoter detection, and transcription factor detection. A recent study has been released that provides a baseline framework for benchmarking these models [20].

Herein, we integrate 16S rRNA gene analysis and embeddings from Genomic Language Models to develop innovative and alignment-free bacterial classification methods. Using existing 16S sequence datasets and sequence embeddings of pretrained gLMs, we assess their potential and limitations in achieving reliable and scalable taxonomic assignments. This study demonstrates that embedding-based sequence classification can match and even outperform BLAST+ [21, 22] and VSEARCH [23] performance in bacterial taxonomy classification at a fraction of computational requirements and help identify novel or mislabeled sequences. Additionally, we highlight key challenges of current sequence embeddings, particularly their sensitivity to sequence length variations. Addressing these limitations is crucial for optimizing gLM applications for microbial taxonomy classification and enhancing their role in metagenomic workflows. Through our findings, our objective is to contribute to the development of more robust, accurate and computationally efficient tools for bacterial classification.

## 2 Results

### 2.1 Phylogenetic groups in the embedding space

We evaluated the effect of the DNABERT-S contrastive training procedure by selecting 10,000 pairs of random sequence embeddings per taxonomic rank from the Greengenes2 [24] dataset belonging to the same phylogenetic group at the same rank (phylum to species) and calculated their cosine similarity. While DNABERT-S efficiently translates natural sequence diversity within the 16S gene to a wide spread of similarity scores (0.2 to 1.0), the intra-phylogenetic rank cosine similarities generated by the DNABERT-2 model are constrained between 0.97-1.0 for 95% of the embedding pairs analysed and between 0.94-1.0 for 95% of the embeddings generated by the NT2 500M model. In contrast, embeddings generated with the Evo 7B model demonstrated 0.999 similarity or higher across all phylogenetic ranks making it barely possible to differentiate larger phylogenetic groups from each other (Figure 1). To visualize the ability of each evaluated model to distinguish sequences from different phylogenetic groups, we used UMAP [25] to calculate a two-dimensional representation of 10,389 sequence embeddings for 20 families of the order Actinomycetota with the highest number of unique database entries. The UMAP projections of the embedding spaces of both DNABERT models showed a remarkable similarity with well-defined family clusters (Figure 2), although DNABERT-2 supports stronger isolation between clusters of phylogenetic groups. In contrast, NT and Evo could not separate taxonomic groups into well-defined clusters. These results indicated that neither Evo nor NT2 is suitable for the 16S sequence classification.

**Figure 1.**
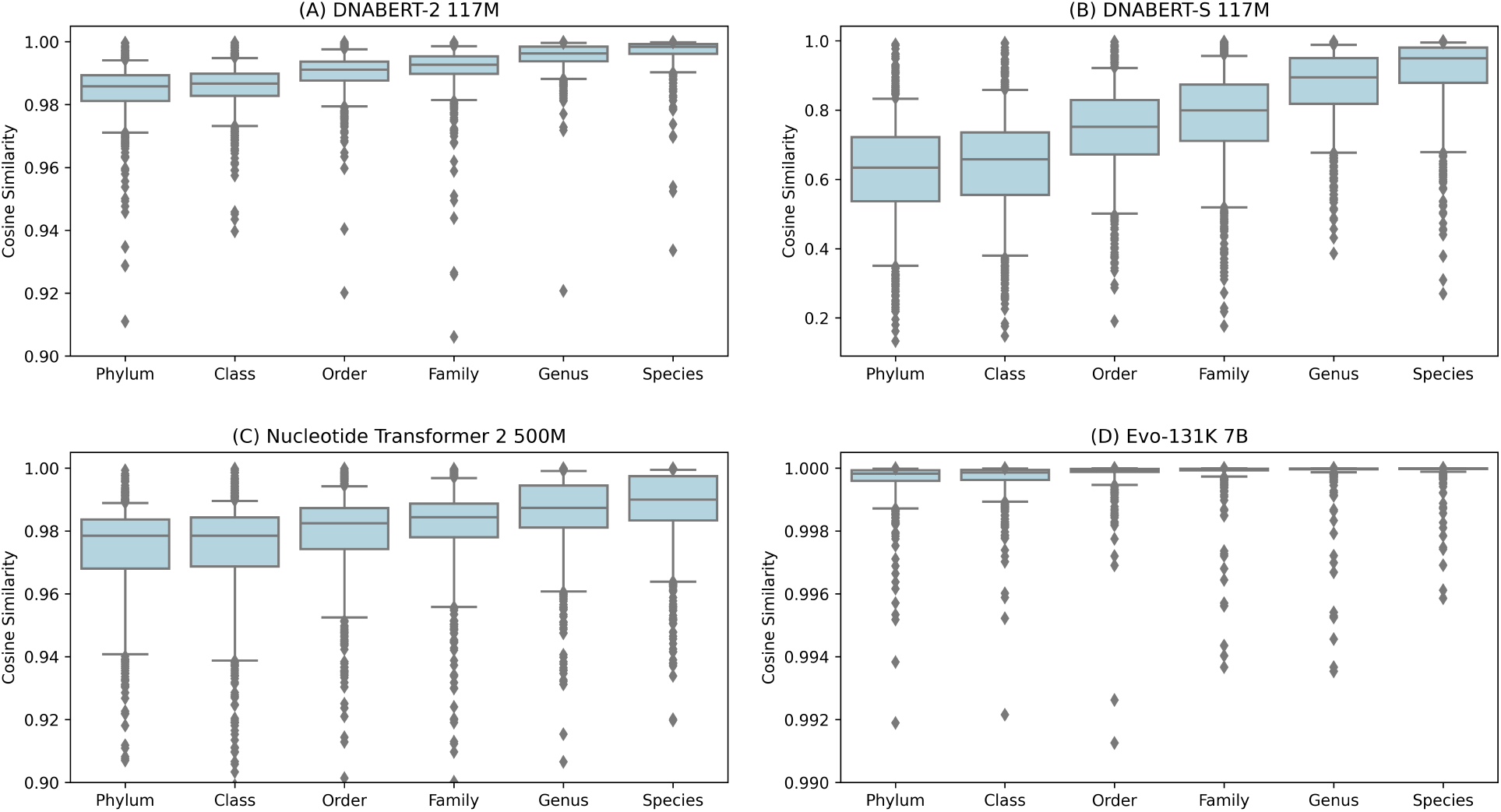
Distribution of cosine similarities of randomly selected sequence embeddings belonging to the same phylogenetic group at the respective taxonomic rank for DNABERT-2 (A), DNABERT-S (B), Nucleotide Transformer 2 500B, and Evo-131K 7B. The average cosine similarity for DNABERT-S was 0.91 for Species, 0.87 for Genus, 0.78 for Family, 0.74 for Order, 0.64 for Class, and 0.62 for Phylum. In contrast, the average cosine similarity for all taxonomic ranks was above 0.97 for DNABERT-2, between 0.94 and 0.99 for NT2, and above 0.999 for Evo 7B. This difference was due to the contrastive loss function used for DNABERT-S finetuning.

**Figure 2.**
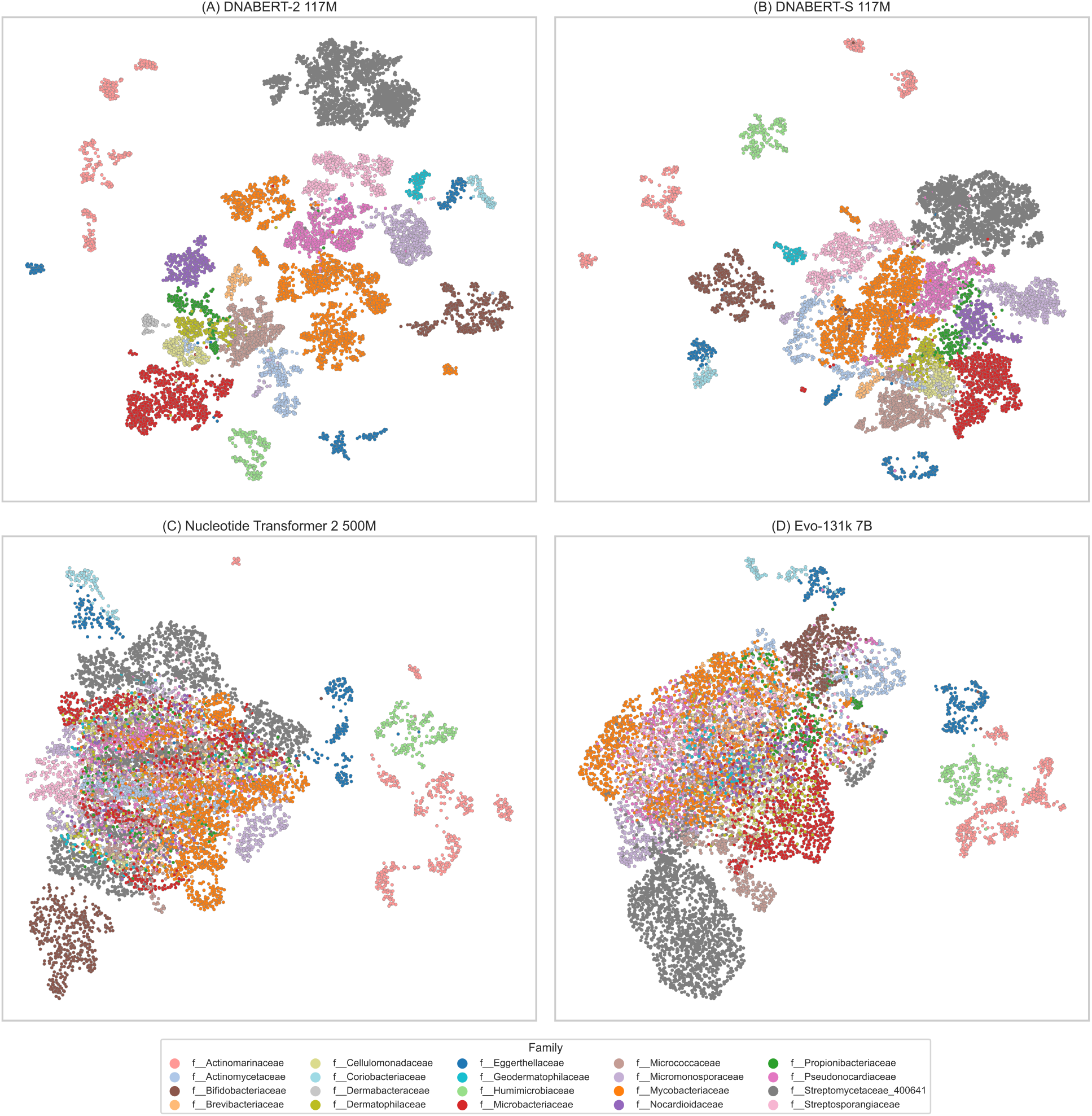
UMAP two-dimensional visualisation of unique 16S sequence embeddings (n=10,689) of individuals belonging to the Actinomycetota phylum, retrieved from Greengenes2 dataset and color coded using family information. Sequences have been embedded with 4 different models: (A) DNABERT-2, (B) DNABERT-S, (C) Nucleotide Transformer 2 500M, (D) Evo-131K 7B. The top 20 families with the highest number of unique database entries are shown to improve visibility.

### 2.2 Embedding-based Classification

Central to this research, we evaluated the ability of various gLM embeddings to provide accurate taxonomic classification of bacterial sequences derived from the 16S ribosomal sub-unit gene. To achieve this, we used the Greengenes2 [24] and GTDB [26] datasets, and focused especially on full-length 16S sequences, as well as on the V3-V4 regions. For each classification task (Greengenes2/Full16S, GTDB/Full16S, Greengenes2/V3V4, GTDB7/V3V4), we present the results using BLAST+ 2.16.0, VSEARCH 2.30.0, default gLM embeddings, and the best-performing dimensionality-reduced embeddings (UMAP down-projected) for DNABERT and NT2 models (Supplementary Table 1). We use a “model@dimension” notation to refer to embeddings from a given model with a particular dimensionality. Although BLAST+ (97.6, 93.9, 90.3, 75.8) outperformed all default model embeddings in classification accuracy, DNABERT-2@768 (97.1, 92.7, 87.7, 72.2) and DNABERT-S@768 (96.2, 91.8, 84.6, 71.1) achieved competitive scores, with the advantage of completing classification in orders of magnitude faster using the embedding-based vector similarity search.

**Table 1:** Common 16S rRNA reference datasets and numbers of groups per taxonomic level. (*) For the Silva dataset, the species number presented here ignores strain information.

| Dataset | Version | File | Phyla | Classes | Orders | Families | Genera | Species |
| --- | --- | --- | --- | --- | --- | --- | --- | --- |
| Greengenes2 [24] | 2024.09 | Backbone full-length | 131 | 334 | 958 | 2,108 | 8,115 | 22,902 |
|  |  | All sequences | 131 | 334 | 958 | 2,108 | 8,115 | 22,902 |
| GTDB [26] | R220 | SSU All | 174 | 468 | 1,584 | 3,978 | 16,872 | 63,859 |
|  |  | SSU Representative | 174 | 458 | 1,553 | 3,884 | 16,117 | 58,102 |
| Silva [37] | 138.2 | SSURef | 97 | 202 | 516 | 755 | 3,857 | 39,357* |
|  |  | SSURef NR99 | 97 | 202 | 516 | 755 | 3,857 | 25,051* |
| RefSeq [41] |  | TargetedLoci/Bacteria |  |  |  |  |  | 19,862 |

In contrast, classifying query sequences using the NT2 500M (90.8, 85.9, 83.1, 64.0) and Evo (87.0, 87.6, 76.7, 64.5) embeddings resulted in a reduced performance in accordance with their limited capability to separate phylogenetic groups as shown in the 2d UMAP visualization of the Actinomycetota embeddings (Figure1). The taxonomy assignment using VSEARCH (97.0, 87.6, 93.9, 73.9) led to accuracy scores marginally lower than DNABERT-2@768 for both Greengenes2 classification tasks, while scoring slightly higher on the GTDB tasks.

Based on these initial classification results, we used UMAP to transform the embeddings of the DNABERT-2, DNABERT-S, and NT2 models into a set ranging from 8 to 256 dimensions, which outperformed the accuracy of the BLAST + classification for all four classification tasks (Figure 3). For the Greengenes2/Full16S classification task, DNABERT-2@128 achieved the best embedding-based classification score with an average accuracy of 98.0, while DNABERT-S@128 achieved a total accuracy of 97.2. BLAST+ and VSEARCH achieved classification scores of 97.6 and 97.0, respectively. On a more granular level, DNABERT-2@128 achieved a higher classification score at species level (DNABERT-2: 96.8, BLAST+: 95.6), while BLAST+ achieved slightly higher scores at genus level accuracy (DNABERT-2: 99.1, BLAST+: 99.5). On the GTDB/Full16S task, DNABERT-2@128 outperformed BLAST+ and VSEARCH by an average accuracy of 2.5 percentage points (DNABERT2: 96.4, BLAST+: 93.9, VSEARCH: 93.9).

**Figure 3.**
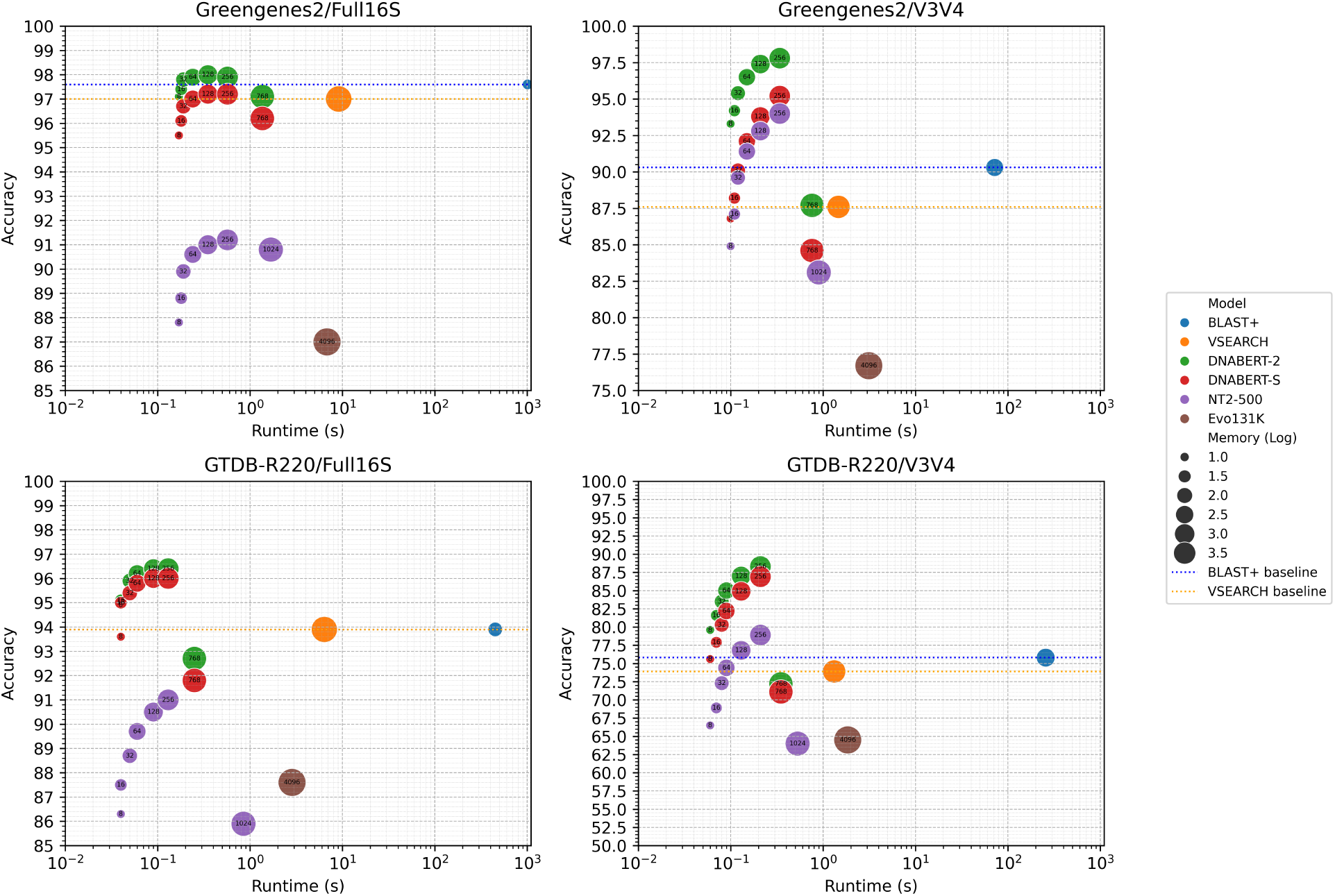
Performance results after employing UMAP to reduce original embeddings to subsets thereof across four datasets leading to up to 11,000-fold reduction in classification time over BLAST+, while outperforming BLAST+ accuracy. Runtime and memory footprint (for the whole dataset) are visualized in log-scale. Memory footprint for BLAST+ reports RAM usage, which was up to 18x less than required for BLAST database. All results collected on an AMD Ryzen 7 5800H with 3.2GHz (16 threads) and 16GB RAM. The measured runtimes do not include the time required to embed the 1,000 query sequences, which takes approximately 8 seconds on a NVIDIA H100 GPU and 12 seconds on a NVIDIA RTX4080 GPU.

For classification using the V3-V4 region, all dimensionality-reduced embeddings achieved higher accuracy scores than the respective model’s default embeddings (Figure 3), with 256 UMAP dimensions achieving the highest performance improvements (DNABERT-2: +10.1/+16.3 (Greengenes2/V3V4 / GTDB/V3V4) percentage points, DNABERT-S: +10.6/+15.8 percentage points, NT2 500M: +10.9/+14.9 percentage points). DNABERT-2@256 achieved the highest classification scores for the Greengenes2/V3V4 (DNABERT-2: 97.8, BLAST+: 90.3, VSEARCH: 87.6) and the GTDB/V3V4 tasks (DNABERT-2: 88.3, BLAST+ 75.8, VSEARCH: 73.9), while DNABERT-S@256 achieved slightly lower accuracy scores. Further details on accuracy results, runtime and memory consumption are provided in Supplementary Table 2. Of the 4 models tested, DNABERT-2 and its down projections proved to be the most accurate regarding the evaluated datasets.

**Table 2:** Overview of Genome Language Models used in this study.

| Model | Model Size | Architecture | Tokenization | Embeddings Size |
| --- | --- | --- | --- | --- |
| **DNABERT-2** [7] | 117M | Bert [10] | BPE [42] | 768 |
| **DNABERT-S** [19] | 117M | Bert [10] | BPE [42] | 768 |
| **NT 2 500M** [6] | 500M | Roberta [43] | 6-mer | 1024 |
| **Evo 131K 7B** [5] | 7B | StripedHyena [44] | ByteCharacter [30] | 4096 |

### 2.3 Sequence Mislabeling Detection

In the course of the analysis, numerous instances were identified in which sequences of different taxonomic groups clustered together, highlighting potentially mislabelled sequences. We identified a total of 4,802 potentially mis-labeled unique 16S sequence-taxonomy entries out of 127,492 unique sequences, with sequence lengths ranging between 1,450bp and 1,550bp in GTDB R220. After removing these potentially mislabeled sequences from the dataset and resolving them against the remaining sequences using the DNABERT-S model, we reclassified 1,916 sequences into taxonomic groups using a cosine similarity threshold greater than 0.95 (DNABERT-S embeddings). For 1,713 sequences, our results suggest a taxonomic mislabeling at the genus level or broader. Noticeably, *Wolbachia, Escherichia, Klebsiella*, and *Staphylococcus* were the genera most frequently affected (Figure 4 (a)). Among the potentially 132 mislabeled *Wolbachia* sequences, DNABERT-S classified 77 sequences as *Acetobacter* and 14 as *Escherichia*; 57 out of these 77 *Wolbachia* sequences have exact duplicate entries in GTDB correctly labeled as *Acetobacter* or *Escherichia*. We further investigated the GTDB sequence annotations for *Staphylococcus aureus*, which is known to cause a wide range of diseases, including skin infections, sepsis, pneumonia, and endocarditis [27]. Pairwise cosine similarity scores within a phylogenetic group of interest quickly identified intra-group classification inconsistencies. Figure 4 (b) highlights 4 outliers among 40 randomly selected Staphylococcus aureus entries in GTDB R220 (RS_GCF_010571055.1 NZ_JAAGQU010000043.1, RS_GCF_027272745.1 NZ_JAPVOH010000020.1, RS_GCF_027944475.1 NZ_JAQJBQ010000008.1, and RS_GCF_010571025.1 NZ_JAAGQW010000067.19), which we classified as *Salmonella enterica* (cosine similarities: 1.0, 1.0), *Escherichia coli* (cosine similarity: 1.0), and *Acinetobacter fasciculus* (cosine similarity: 0.9959). These classifications were confirmed by NCBI Web BLAST (May 2025, core_nt database) except for *A. lwoffii* instead of *A. fasciculus*. The 2d UMAP projection of the sequence embeddings of the respective species confirms the identified sequence misannotations and raises further quality concerns, as additional mutual misclassifications between the 5 classes are visible alongside two clusters of sequences for which the true identity is not covered by this UMAP projection (Figure 4 (c)). We provide a list of 24 probably mislabeled Staphylococcus aureus GTDB entries in Supplementary Table 3.

**Figure 4.**
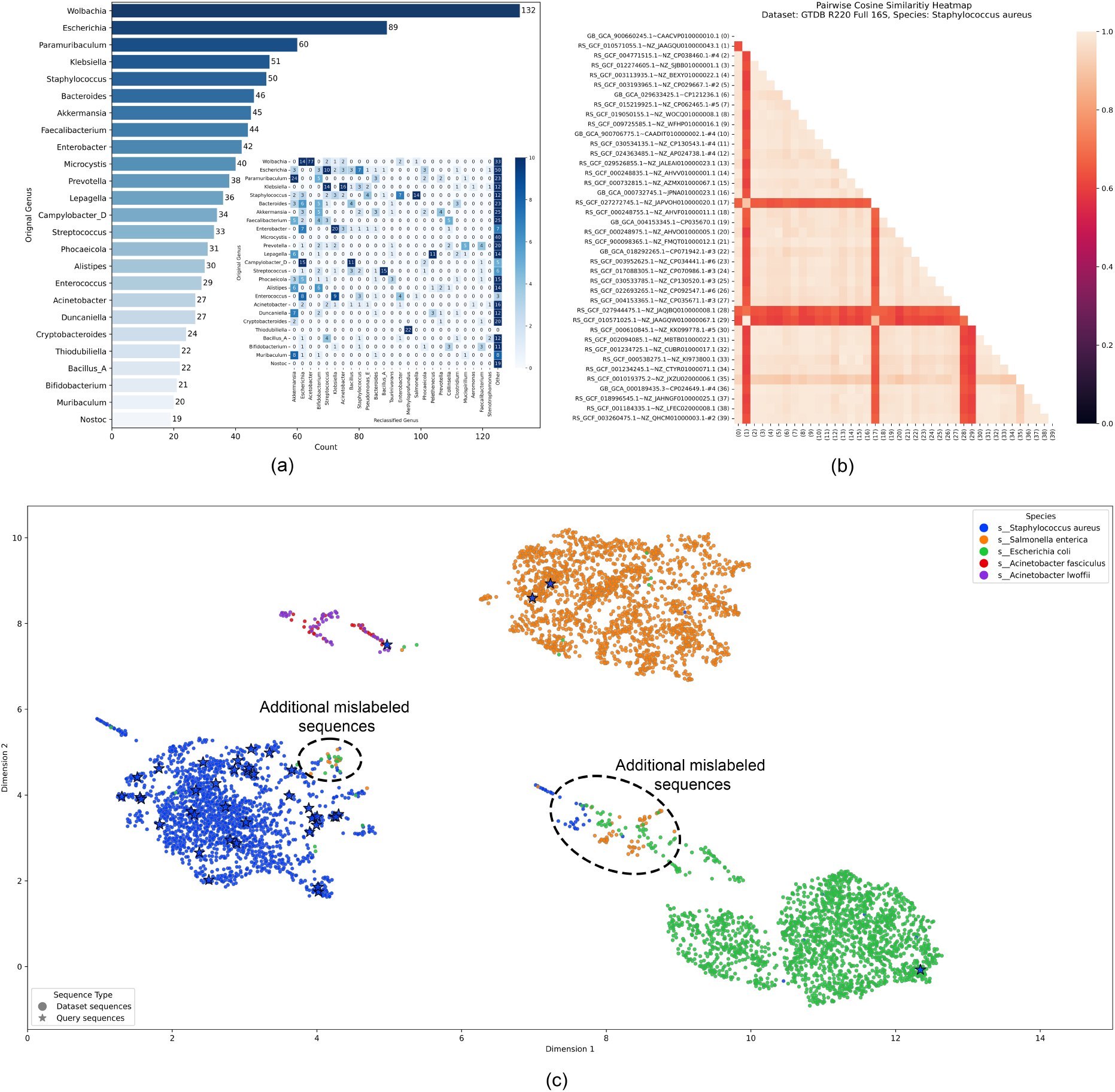
Mislabeled sequence identification and reclassification workflow for GTDB R220. (a) Overview of the 25 most frequently mislabeled genera and heatmap visualization of mutual mislabeling among the 25 most frequent genera. Matching genera indicate a proposed taxonomy misannotation on the species level. (b) Outlier detection using pairwise cosine similarity computation quickly highlights inconsistencies within a phylogenetic group. We randomly selected 40 S. aureus sequences for this analysis. (c) 2d UMAP projection of the respective species, where stars represent 40 entries from (b). The number of sequences per species was capped at 2,000 for improved visualization. (a-c) All embeddings were generated using the DNABERT-S model.

In contrast to the GTDB results, the same analysis on the Greengenes2 full-length backbone dataset resulted in only 110 potentially mislabeled sequences (after removing sequences with unnamed species), of which 35 sequences indicate a different genus after reclassification.

### 2.4 Sequence-length Dependency

DNABERT-S embeddings of subsequences (1,400bp long) extracted from a full 16S sequence (~1,500bp) will, on average, lead to a cosine similarity of approximately 0.95 to its original sequence, making it often difficult to identify its correct full-length taxonomy (Figure 5). For sequences that have been shortened by 200 base pairs, the average cosine similarity to the full-length sequence fell below 0.9. A similar degradation of similarity scores was observed with the embeddings generated by DNABERT-2, albeit proportionate to the smaller range of similarity values (Figure 3). The sensitivity shown by these models to truncation of sequences is a stark contrast to their performance on the V3-V4 segment, which, despite being only ~400bp long, achieved strong classification results when compared against other V3-V4 embeddings.

**Figure 5.**
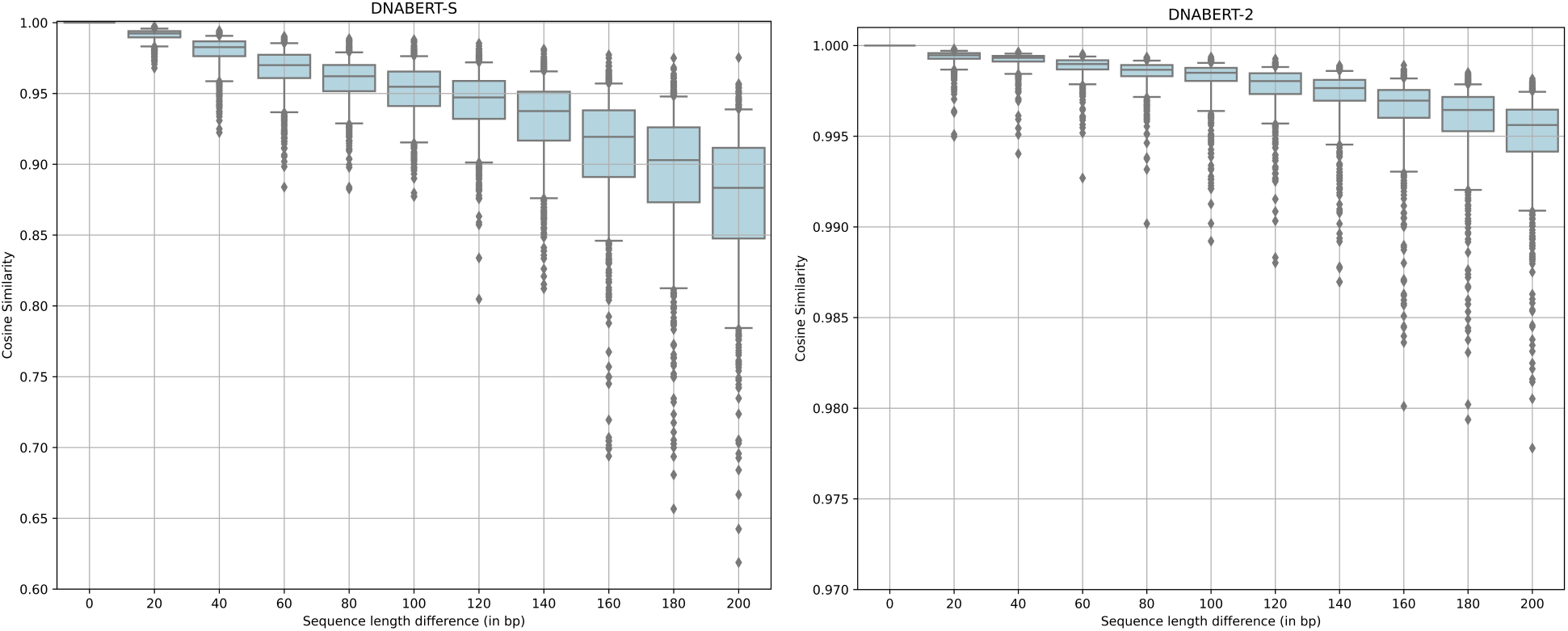
Decline of average cosine similarity between the original sequence embeddings and embeddings of iteratively shortened sequences. Box represents mean similarity, together with 25% and 75% quartiles. Whiskers represent 5% and 95% quantiles, respectively.

## 3 Discussion

Taxonomic classification is a central step in metagenomic workflows because it allows the identification and categorization of microbial DNA reads present in a sample. The assignment of sequences to known taxonomic groups helps interpret microbial diversity, community structure, and functional potential within an environment. It provides a foundation for comparing microbial populations across different conditions (e.g., healthy versus diseased states in humans) and identifying specific microbes associated with certain ecological roles or health outcomes. Alignment-based methods have been the gold standard for microbial taxonomic classification for the past six decades, despite their computational expense and slow processing speeds. Although alignment-free approaches have been developed to address these limitations [28, 29], many still suffer from similar performance drawbacks. In this study, we demonstrate the potential of embedding-based methods for 16S rRNA gene classification. We evaluate multiple methodologies, with a particular focus on embedding-based classification, and compare their performance, including CPU time, memory usage, and precision, to gold-standard tools such as BLAST and VSEARCH.

### 3.1 Token-based classification versus embedding-based classification

The success of GPT and LLaMA-based models has set expectations for language models to generate tailored responses to given queries. This capability is driven by the decoder component of language models, which leverages the internal representation of an input sequence to predict the most probable next token. In the context of Genomic Language Models (gLMs), this predictive mechanism can be naturally applied to nucleotide sequences, forecasting the next nucleotide(s) in a given sequence. In token-based taxonomy classification approaches, the gLM requires a model vocabulary that extends beyond nucleic bases towards a complete alphabet, as well as special characters included in taxonomic naming.

The Evo model uses a ByteCharacter tokenizer [30] that converts 255 ASCII characters into tokens, allowing the integration of nucleotide sequences with natural language, such as the taxonomic label of a sequence. By inverting a part of Evo’s pre-training in such a way that the model returns natural text given an input DNA sequence, the model can infer taxonomy directly from the input sequence. However, being a generative language model, Evo is not suitable for taxonomy classification since results can be fragile: even minor output sequence errors, like a single misspelled character, will lead to invalid taxonomy strings such as “Escherichia cali” that would need to be caught using fuzzy matching.

Introducing special taxonomy-related tokens poses a simplified form of the iterative next-token classification in which only a single output token needs to be generated; for example, a species-specific token. However, introducing tens of thousands of species-level tokens into a model’s vocabulary requires extensive model training to effectively learn the often nuanced relationships between a sequence and the correct bacterial species. In addition, both token-based classification approaches face a severe limitation in the model’s ignorance of rearrangements or additions to the taxonomic landscape, and its inability to adapt to databases with different naming conventions without repeated model fine-tuning.

For gLMs, embeddings are vectorized representations of DNA sequences, where similar sequences produce similar vector representations, which can then be used for downstream analysis and classification tasks. Using known sequence-taxonomy assignments from curated 16S datasets, classical machine learning algorithms or neural networks can be trained using the supervised learning paradigm to learn the relationship between a set of sequence embeddings and the corresponding taxonomy labels. However, this leads to the same fundamental algorithmic difficulties as described for taxonomic classification with tokens: the model must learn to differentiate between tens of thousands of classes for which typically only a few samples per class are available during the training process, as well as the requirement for repeated model training when reference databases are updated.

Unlike algorithms that train a classifier on sequence embeddings, *similarity search* directly compares embeddings to existing labeled sequences without explicit model training. Instead of learning a decision boundary, it finds the most similar embeddings in a vector store populated with precomputed sequence embeddings. That makes similarity search achievable in scenarios with few samples per class, real-time retrieval, and handling new or evolving categories without retraining a model. A commonly used similarity metric is the *cosine similarity*, representing the similarity of two vectors regardless of their size, and is calculated as the dot product between two vectors divided by the product of the vector lengths, being mathematically simple and fast, irrespective of the number of dimensions analyzed.

### 3.2 Embedding-based search and classification accuracy

While decoder-only models (i.e., Evo) are primarily designed to predict the most likely continuation of a sequence and are not explicitly optimized to capture biologically meaningful differences between sequences, encoder-only models (i.e., DNABERT-2 and Nucleotide Transformer) are generally suitable for classification tasks. Both DNABERT models demonstrated that embedding-based taxonomic classification of 16S sequences can lead to computationally efficient and accurate novel bioinformatic workflows. The main difference between these models is that DNABERT-S was trained using a contrastive loss function (based on cosine similarity) to ensure that nearly identical DNA sequences produce similar vector representations while spreading dissimilar sequences farther apart in the vector space. This model training paradigm led to worse accuracy scores due to less defined decision boundaries between phylogenetic groups when applied to the 16S gene. Using the default DNABERT-2@768 model embeddings resulted in competitive results compared to BLAST+, although it required two to three orders of magnitude less computational runtime (0.25s compared to 451s for BLAST+ and 6.37s for VSEARCH regarding the GTDB/Full16S dataset) (Supplementary Table 2).

By reducing the dimensionality of the embedding space from 768 to between 8 and 256 dimensions, we improved not only the runtime and memory usage but also the accuracy itself. In fact, the UMAP-based reduction in dimensionality of the gLM embeddings is transformative, enabling us to match or surpass the accuracy of the BLAST+ (and VSEARCH) results across all classification tasks. This method increases taxonomic accuracy to ~97% for species, and more than 99% for broader taxonomic levels such as genus and family when using embeddings generated using Greengenes2 datasets. The largest performance leaps were observed for the classification of the V3-V4 sequence, where DNABERT-2 surpassed BLAST+ by up to 12.5 percentage points, suggesting that shorter sequences do not need a 768-dimensional space for biologically meaningful representations and could benefit more than longer sequences from restricting the vector space (Supplementary Table 1). This improvement in species-level classification beyond the state-of-the-art for V3-V4 is especially impactful given that microbiology relies heavily on 16S subsequences (e.g., V3-V4) for identification, despite the limited reliability in classifying subsequences at species level via classical approaches [31–34].

Furthermore, DNABERT-2 embeddings outperformed DNABERT-S embeddings in each experiment by up to 1.6 percentage points in full-length sequences and up to 6.5 percentage points for V3-V4 sequences. In total, sequence classification against the Greengenes2 dataset led to higher accuracy scores than against the GTDB dataset. We expect this to be grounded in the overall larger number of sequences included in the former dataset and the surprisingly high number of mislabeled sequences included in the GTDB dataset (Supplementary Table 1).

### 3.3 Dimensionality reduction of the embedding space improved runtime and memory usage

In addition to classification accuracy, runtime performance and memory usage are critical computational metrics for evaluating sequence analysis methods. Transforming DNA sequences from their raw string format into fixed-length numerical vector representations enables the use of highly optimized linear algebra operations for downstream analysis. In particular, sequence similarity computations can be performed efficiently via matrix multiplication, a fundamental operation extensively optimized on both modern CPUs and GPUs. Matrix multiplication is computationally efficient due to its regular memory access patterns and suitability for parallelisation. Typically, each matrix element is stored as a 32-bit floating point number (4 bytes), and the number of arithmetic operations involved in matrix multiplication scales with the dimensionality of the vectors: multiplying two matrices of shapes NxD and DxM requires O(NxDxM) operations. As such, higher-dimensional embeddings result in increased computational cost, underscoring the importance of balancing representational richness with processing efficiency.

A key advantage of an embedding-based search lies in its ability to increase classification performance even when using substantially reduced embedding dimensions, allowing us to reduce classification runtime by an order of magnitude. For the Greengenes2/Full16S task, DNABERT-2@128 achieved the highest classification score in 0.35s, while it took 9.12s for VSEARCH and 1010s for BLAST+. DNABERT-2@32 achieved a marginally lower score in 0.19s, still outperforming BLAST+. DNABERT-2@256 achieved the highest classification scores for the other 3 classification tasks (GTDB/Full16S, Greengenes2/V3V4, GTDB/V3V4) in 0.13s, 0.34s, and 0.21s, respectively. For these tasks, even DNABERT-2@8 outperformed the established tools in 0.04s, 0.10s, and 0.06s. Contrarily, VSEARCH completed these tasks in 1.47s, 6.37s, and 1.32s. BLAST+ demanded the largest computational resources with runtimes of 451s, 72s, and 257s. These findings underscore the significant computational paradigm change that embedding-based classification will impose on future bioinformatic workflows. Further optimizations, such as vector quantisation or hierarchical indices, were not evaluated for the vector-based search.

Besides the runtime improvements, the reduced projection of the embeddings into a lower dimensionality led to a reduction in memory footprint, which decreases linearly following a simple paradigm: *number of sequences * number of dimensions * 4 bytes per dimension*. Depending on the number of sequences, the default DNABERT embeddings required between 238 MB (GTDB/V3V4: 80,059 sequences) and 861 MB (Greengenes2/Full16S: 293,642 sequences). Using the best-performing down-projected embeddings (128, 256) decreased the memory footprint to 80 MB to 144 MB. With the smallest number of dimensions (8), the embedding matrix required between 3 and 9 MB. Across the 4 classification tasks, BLAST+ required between 53 MB and 206 MB disk space for the database and 12-40 MB RAM usage. VSEARCH required up to 2,100 MB for the Full16S datasets and up to 845 MB for a V3-V4 dataset.

### 3.4 Embedding similarity metric effectively identifies mislabeled sequences

Taxonomic mislabelling (or misannotation) refers to the incorrect assignment of taxonomic identity to a sequence, often resulting from errors in sequencing or taxonomic identification by data submitters [35]. According to Chrolton [2], this widespread issue stems from a lack of standardized taxonomic identification protocols, compounded by inconsistencies in reference database taxonomies - such as those in RDP [36], SILVA [37], and Greengenes, each of which employs distinct classification frameworks [38, 39] [40, 41]. Recent efforts to address misannotation include initiatives like the Genome Taxonomy Database (GTDB) and Greengenes2, which integrate microbial data from 16S rRNA gene sequencing and shotgun metagenomics using unified methodologies [24, 40].

We used the DNABERT-S model to investigate the impact of mislabeled sequences within the two databases due to its training regime, where the use of a contrastive loss function generated cosine similarities with well-defined ranges for different taxonomic ranks (Figure 2). Based on previous results (Figure 2), we used a threshold of 0.9 for intra-species cosine similarity and identified 3.77% (4,802) of GTDB’s unique full-length sequences (1,450-1,550bp) as potentially mislabeled. Of these, 1,916 sequences showed cosine similarity scores between 0.95 and 1.0 against other known taxonomic groups from GTDB R220, strongly suggesting their true taxonomic identity as different from their original classification. For 1,713 sequences, our reclassification results suggest a sequence mislabeling on the genus level or broader. Although several well-studied genera such as *Klebsiella, Wolbachia*, and *Escherichia* have been identified as being mislabeled especially frequently, more than 440 genera in total are affected in the GTDB dataset, underlining the potential for distortion of microbiological studies. More than 96% of these mislabeled sequences have been identified as metagenome-assembled genomes (MAG) in NCBI, highlighting the importance of quality control for any future secondary rRNA 16S dataset curated from NCBI sources. In contrast, using the same procedure on the Greengenes2 full-length backbone dataset filtered to retain fully-named sequences resulted in a limited number of 35 mislabelled sequences, where the embedding-based classification indicates a misannotation on genus level or broader taxonomic ranks.

### 3.5 Limitations of Embedding-based models for DNA sequence classification

Despite their accuracy and computational advantages over established methods, embedding-based sequence classification faces key limitations that can hinder its practical application. The components of the rRNA operon (5S, 16S, 23S) are characterized by their high information density, making them ideal for taxonomic classification; however, this same attribute poses challenges for sequence embeddings, as variations in the query sequence length can lead to significantly different embeddings. As a result, even minor length differences between two otherwise identical sequences can result in embeddings that fail to capture their biological similarity. Ideally, sequence embeddings should produce similar vector representations for sequences that are subsets of each other. For example, a full-length 16S sequence and the V3-V4 region of the same sequence should yield comparable embeddings. In practice, however, length differences, such as a 1,500bp sequence compared to a 1,450bp subsequence, can result in embeddings that differ considerably, impacting the classification of shorter sequences.

The results outlined in the section “Sequence-length Dependency” imply that the current sequence embedding comparison is similar to the traditional Global Alignment methods, where all sequences are expected to be of the same length, rather than Local Alignment, which allows the association of shorter sequences against longer ones. Further research into length-insensitive embeddings (or model architectures) is required to achieve a classification comparable to Local Alignment in vector space. To address the length sensitivity short term, systems using embedding-based taxonomy classification can precompute embeddings for common subsequences, such as those corresponding to specific hypervariable regions or full genes with similar gene sizes, such as the 16S rRNA coding gene. However, these systems will struggle with random reads of shotgun sequences, which lack region information. In contrast, traditional alignment-based classification systems like BLAST are inherently length-independent. They can reliably match subsequences of any length to their corresponding full-length sequences database, providing a robust baseline for classifying sequences with variable lengths.

Another limitation is the lack of standardized similarity thresholds, as different gLMs produce varying inter-and intra-taxonomic-group similarity scores, impacting the establishment of universal classification criteria. Likewise, establishing a new benchmark dataset for embedding-based taxonomy classification will help evaluate the impact of future enhancements in gLM architectures or training methods on classification accuracies and the current limitations identified in this study. Token-based classification methods were excluded from evaluation as it is evident that they represent no scalable future-proven approach due to their inability to identify novel sequences and their need for model retraining when new phylogenetic classes are discovered and assigned.

## 4 Conclusion

This study explored the adoption of Genomic Language Model embeddings for bacterial taxonomy classification using 16S rRNA sequences. Transformation of sequences to numerical vectors allows efficient vector operations to calculate sequence similarity, outperforming traditional alignment-based tools like BLAST+ and VSEARCH in classification accuracy, while completing the classification process up to 4 orders of magnitude faster. DNABERT-2 achieved the highest overall accuracy scores in our experiments, and its embeddings seemed well suited for the taxonomy classification of bacterial sequences. UMAP-based embedding projections not only led to a lower dimensionality of the numerical representations of DNA sequences but are a cornerstone for achieving state-of-the-art classification performance while optimizing system-level resources such as runtime and memory footprint. In addition to the classification performance, the sequence embeddings demonstrated the ability to efficiently detect mislabeled sequences. These results position sequence embeddings as a viable alternative to traditional alignment-based methods, particularly for large-scale applications suffering from slower computation and high memory demands. We further identified a critical limitation in the length sensitivity of embeddings, rooted in how Transformer-based model architectures perform attention calculation across the complete sequence provided, which hinders the classification of sequences with varying lengths. Addressing this limitation offers avenues for further research and advancing the applicability of Genomic Language Models. The rapidly growing volume of metagenomic data demands tools for accurate, efficient, and reliable sequence taxonomy assignments, underscoring the need for advancing methods that represent DNA sequences in computationally efficient forms. Our findings show that gLM-based classification can serve as a scalable and robust alternative to traditional tools, paving the way for fast and high-throughput metagenomic workflows.

## 5 Methods

The central challenges to the taxonomic classification of 16S rRNA sequences from microbes are defined by the large number of potential labels to be predicted. Classifying sequences on the species level, albeit difficult without using the full 16S sequence, requires several tens of thousands of labels; at the genus level, likewise more than 10,000 labels; several thousand labels for individual bacterial families, and so on. Table 1 summarizes the bacterial taxonomy of some of the most commonly used 16S datasets.

### 5.1 Data Acquisition and Preprocessing

The Greengenes2 2024.09 and GTDB R220 SSU data sets were obtained from their respective repositories (Greengenes2, GTDB). For full 16S analysis, both datasets have been preprocessed such that only sequences ranging from 1,450bp to 1,550bp were kept, followed by a data deduplication step to keep only a single entry per sequence-taxonomy pair. This resulted in 293,642 unique sequences (441,174,030 nucleotides) across 18,301 distinct species for Greengenes2 and 127,492 unique sequences (194,284,517 nucleotides) across 37,013 distinct species for GTDB R220. For the V3-V4 analysis, we extracted the corresponding sequence region using the universal primers 341F (S-D-Bact-0341-b-S-17: CCTACGGGNGGCWGCAG) and 785R (S-D-Bact-0785-a-A-21: GACTACHVGGGTATCTAATCC), filtered out all resulting sequences outside the 390bp to 440bp range and applied the same data deduplication step such that per unique sequence-taxonomy pair, only a single embedding was kept. This resulted in 134,186 unique sequences (56,049,976 nucleotides) across 20,494 distinct species for Greengenes2 and 81,059 unique sequences (33,888,309 nucleotides) across 43,821 distinct species for GTDB R220. For each remaining sequence in any dataset, we tracked a count attribute that represents the number of duplicate sequences represented by this sequence. The data deduplication step was performed because identical sequences lead to identical embeddings and, hence, identical points in vector space.

### 5.2 Models

We downloaded the pretrained models of DNABERT-2, DNABERT-S, Nucleotide Transformer 2 500M, and Evo-131k 7B from the HuggingFace Hub (*zhihan1996/DNABERT-2-117M, zhihan1996/DNABERT-S, InstaDeepAI/nucleotide-transformer-v2-500m-multi-species, togethercomputer/evo-1-131k-base*) to generate sequence embeddings. Each model has been trained on multispecies genome data and thus is a prospective candidate for generating robust bacterial sequence embeddings. We employed mean pooling at the last hidden layer to convert the input sequence into an embedding representation. Table 2 highlights the key characteristics of the models used. All of these models were pretrained on multi-species genome data, but only DNABERT-S, a fine-tuned version of DNABERT-2, was trained with a contrastive loss function to spread dissimilar sequences farther apart in the embedding space.

### 5.3 Embedding-based Taxonomy Classification

Computing the cosine similarity and vector search are exceptionally cost-efficient operations on recent processors, allow comparing query sequence embeddings against billions of database entries, and operate multiple orders of magnitude faster than traditional alignment-based approaches. Figure 6 visualizes the overall setup for an embedding-based sequence classification. Initially, the sequences of an rRNA 16S dataset are embedded using a gLM and inserted into a vector store. For this study, we used FAISS [45, 46] (*type*=“Flat”, *metric*=“METRIC_INNER_PRODUCT”) as an efficient in-memory vector store. Subsequently, a user can provide a query sequence for which a bacterial taxonomy needs to be determined for a hypothetical application. The application first embeds the query sequence with the same gLM and then queries the vector store/database to retrieve the *k* closest embeddings to identify the most similar DNA sequences. FAISS search was executed with all 1,000 query sequences as a single batch.

**Figure 6.**
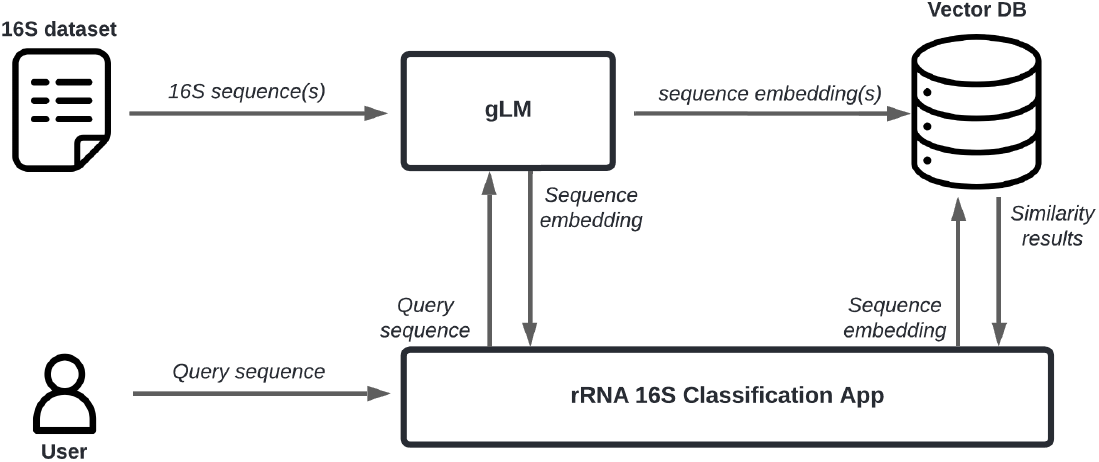
Workflow for embedding-based taxonomy classification

After data acquisition and deduplication, all Greengenes2 and GTDB sequences were embedded using DNABERT-2, DNABERT-S, Nucleotide Transformer 2 (500M), and Evo 131K 7B. For each set of embeddings, we applied dimensionality reduction to reduce the number of dimensions down to two to inspect the embedding space of unique *Actinomycetota* 16S sequences (n=10,689) retrieved from the Greengenes2 dataset. The UMAP library was employed for nonlinear dimensionality reduction to graphically represent sequence embeddings in 2D (parameters: *n_neighbors*=25, *min_dist*=0.75, *metric*=‘braycurtis’, *n_components*=2, *spread*=0.75). UMAP transforms complex datasets into a lower-dimensional space while preserving structural relationships between data points.

To evaluate embedding-based 16S taxonomic classification accuracy, we focused on two targets, the full 16S (1450-1550bp), and V3-V4 region (390-440bp). The UMAP algorithm (parameters: *n_neighbors*=10, *min_dist*=0.25, *metric*=‘braycurtis’, *n_components*=[8, 16, 32, 64, 128, 256]) was applied to project the default embeddings of the DNABERT-2, DNABERT-S, and NT2 models to a range of fewer dimensions, next the appropriate embedding collection was loaded into FAISS, and a total of 100 runs with 1,000 randomly selected query sequences that belonged to named species with at least 2, 3, or 5 sequences were performed. Setting the lower bound for query sequence abundances to at least two was required to still measure classification performance on the species level, such that for each query sequence at least one embedding of the same species resided in the vector store. This experimental setup required 2,100,000 (7 embedding dimensionalities * 3 species selection thresholds * 100 runs * 1,000 queries) sequence classifications per model per classification task. The Evo model was only evaluated using its default embedding resulting in 300,000 total classifications. Due to runtime limitations BLAST+ 2.16.0 [21, 22] was evaluated only against a single run of 1,000 query sequences per species selection threshold per classification task. VSEARCH 2.30.0 [23] performance was evaluated using all 300,000 query sequences (3 species selection thresholds * 100 runs * 1,000 queries) per classification task (parameters: *–usearch_global –id* 0.97 *–qmask* none *–dbmask* none *–top_hits_only –maxaccepts* 10(Full16S)/50(V3V4) *–maxhits* 10(Full16S)/50(V3V4)).

The cosine similarity of the query embeddings was calculated against the appropriate collection using the formulae

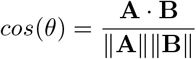

The genus-and species-level taxonomic information of the embedding with the highest similarity score is reported as classification result. For full-length sequence classification, the ten most similar sequences were retrieved from the vector store as in rare occurrences multiple sequences share the Top-1 rank due to sequence mislabeling. For the V3-V4 sequence classification, the 50 most similar sequences were retrieved from the vector store as typically many species of the same genus share a common V3-V4 region and, hence, produce identical embeddings. The accuracy was defined as the proportion of time in which the correct taxonomic label was identified as having the highest similarity score. We compared models and established tools using the average genus and species classification accuracy across the three species selection thresholds. Besides the classification accuracy we tracked the classification runtime and memory consumption to compare the traditional tools BLAST and VSEARCH against the embedding-based classification.

### 5.4 Mislabeled Sequence Detection

The Greengenes2 and GTDB R220 full-length 16S datasets were used in combination with contrastively trained DNABERT-S embeddings to calculate the cosine distances of all unique sequences to the mean embedding of their assigned taxonomic species. Sequences with a cosine similarity of less than 0.9 to their respective average species embedding were flagged as potentially mislabeled. These sequences were then removed from the respective dataset and reclassified against the remaining sequence embeddings to investigate their probable corrected taxonomic identity. We further randomly selected 40 *Staphylococcus aureus* sequences from the GTDB/Full16S dataset and calculated pairwise cosine similarity scores to identify outlier sequences with consistently lower similarity scores. We confirmed the expected true species of those outlier sequences using our embedding-based classification framework as well as NCBI Web BLAST (May 2025, core_nt database). Lastly, we selected up to 2,000 sequences labeled as *S. aureus* including the 40 sequences from the pairwise similarity analysis and from the proposed true species of the outliers and transformed their 768-dimensional embeddings into 2 dimensions using UMAP (*n_neighbors*=7, *min_dist*=0.25, *metric*=‘braycurtis’, *n_components*=2, *spread*=0.3) to visualize the sequence mislabeling.

### 5.5 Sequence Length Dependency

To evaluate the impact of sequence length on vector search, a total of 1,000 random full 16S rRNA sequences were selected from the Greengenes2 dataset. Next, we iteratively removed 10 nucleotides from the beginning and end of the sequence 10 times, leading to stepwise up to 200 base pairs shorter subsequences. For each iteration, we calculated the cosine similarities (using DNABERT-S and DNABERT-2 embeddings) between the original full 16S sequence embedding and their respective shorter subsequences.

## 6 Author Contributions

**Mike Leske:** Conceptualization, Methodology, Formal analysis, Investigation, Visualization, Writing - Original Draft, Writing - Review Editing. **Jamie Fitzgerald:** Formal analysis, Investigation, Visualization, Writing - Original Draft, Writing - Review Editing. **Keith Coughlan:** Investigation, Writing - Original Draft. **Francesca Bottacini:** Methodology, Writing - Original Draft, **Haithem Afli:** Conceptualization, Supervision, Funding acquisition, Writing - Original Draft, **Bruno Gabriel Nascimento Andrade:** Conceptualization, Investigation, Supervision, Funding acquisition, Writing - Original Draft, Writing - Review Editing.

## 8 Supplementary Data

### 8.1 Supplementary Table 1

**Supplementary Table 1.**
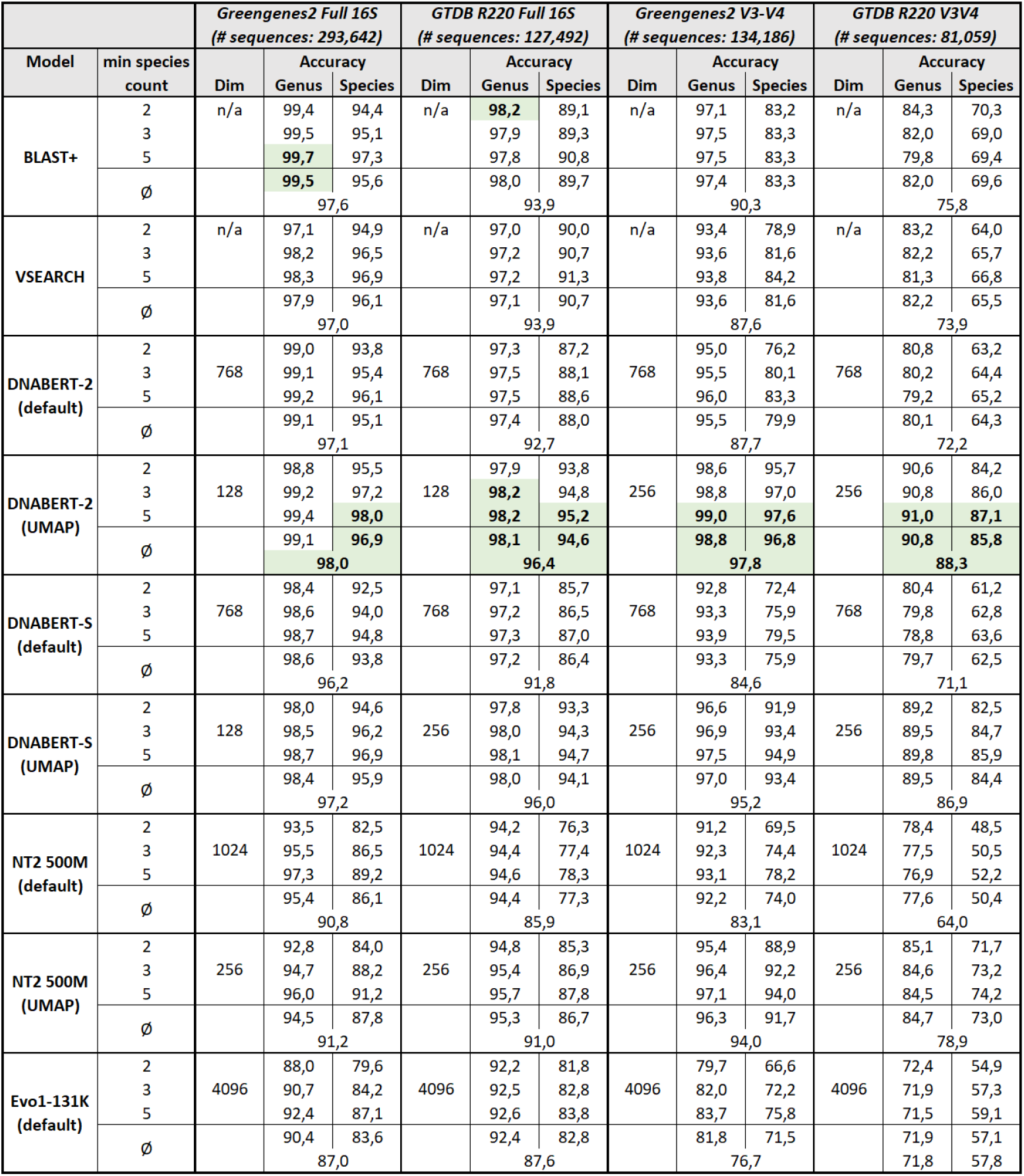
Embedding-based classification results for full 16S rRNA sequence embeddings and V3-V4 subsequence embeddings. Sequence embeddings were generated from the Greengenes2 and GTDB R220 datasets with DNABERT-2 and DNABERT-S models. The table reports the accuracy for genus and species level for the best-performing embeddings (after UMAP dimensionality reduction). Query sequences were sampled from species with at least 2, 3, or 5 distinct sequences. Accuracy represents the fraction of query sequences whose nearest neighbor belonged to the correct phylogenetic group per taxonomic rank. Green cells with bold values represent the highest scores per experiment and accuracy scope (genus, species, average).

### 8.2 Supplementary Table 2

**Supplementary Table 2.**
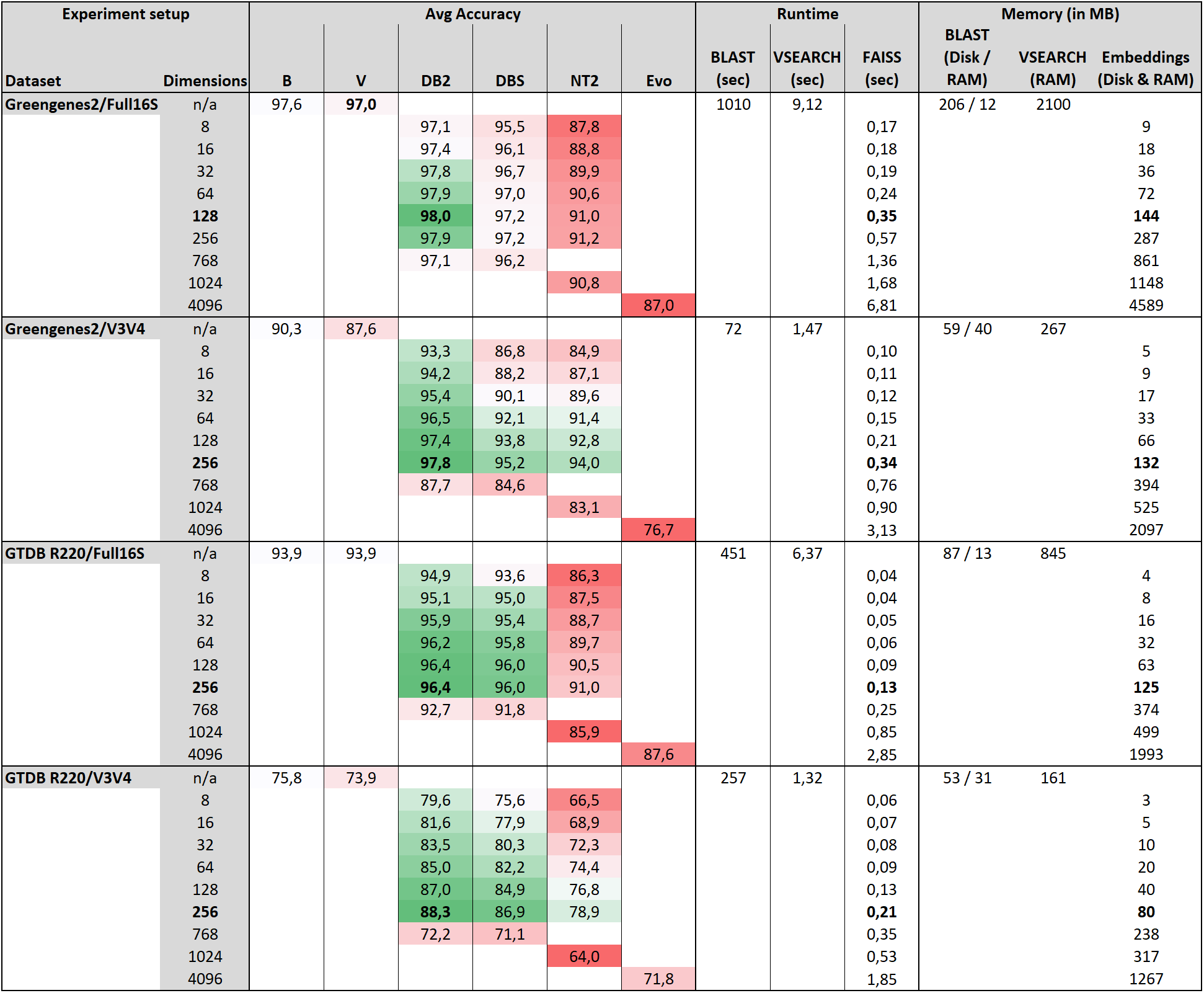
Detailed overview of classification results, runtime, and memory consumption (B=BLAST, V=VSEARCH, DB2=DNABERT-2, DBS=DNABERT-S, NT2=Nucleotide Transformer 2). Bold rows represent the highest classification performance per classification task. Accuracy color gradient with respect to BLAST baseline results (green=higher than BLAST, red=lower than BLAST). Runtime measures for embeddings-based classification measures pure classification runtime and does not include time to embed the 1,000 query sequences. BLAST and VSEARCH runtimes do not include database/index generation.

### 8.3 Supplementary Table 3

**Supplementary Table 3.**
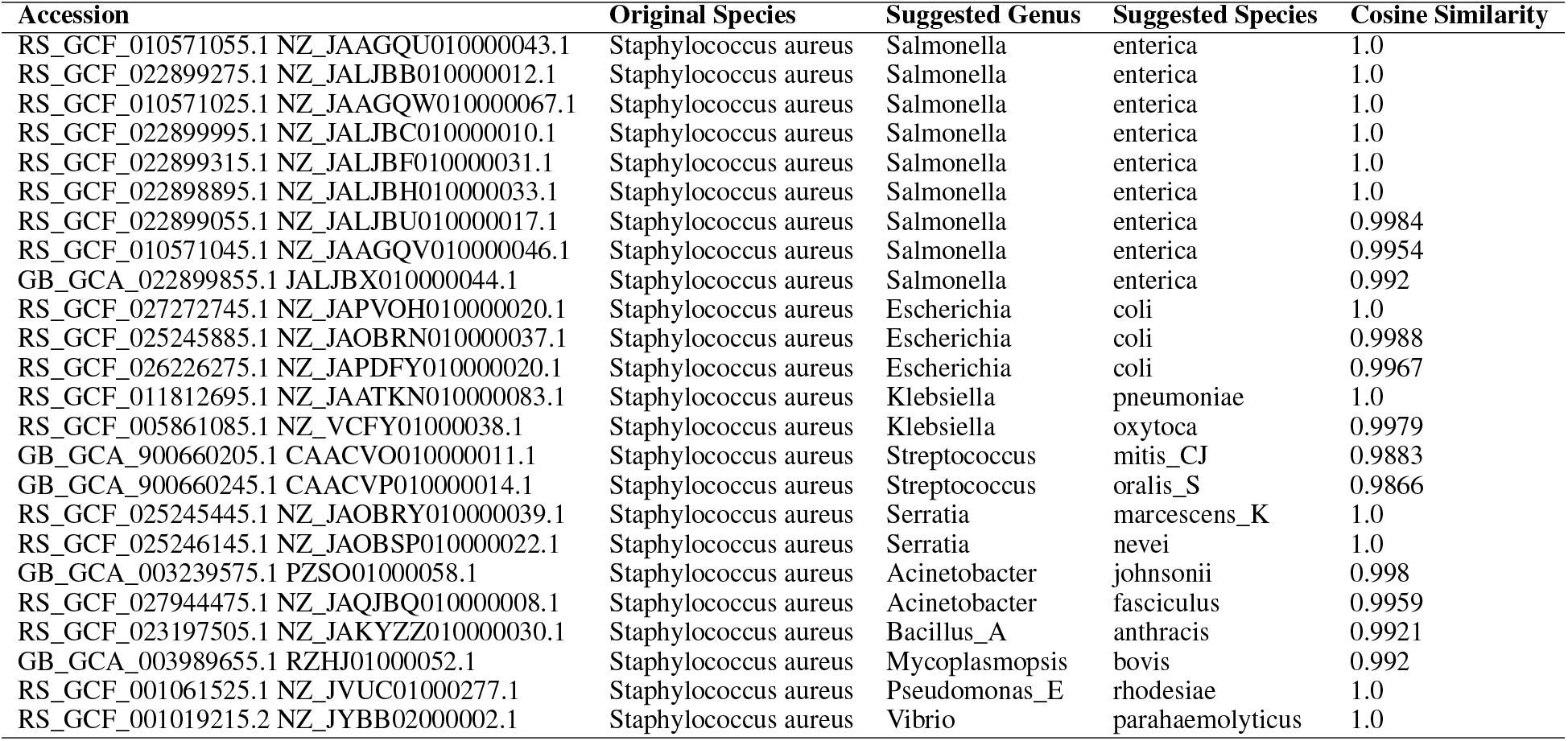
24 potentially mislabeled S. aureus sequences from the GTDB dataset together with a suggested genus and species level reclassification according to our embeddings-based approach. The cosine similarity score represents the reclassification score for the suggested species.

